# Keeping salamanders off the streets: Evaluating one of the first US amphibian road tunnels 30 years later

**DOI:** 10.1101/569426

**Authors:** Brandon P. Hedrick, Abby Vander Linden, Samantha A. Cordero, Edward Watt, Patrick M. O’Roark, Samantha L. Cox, Christopher Sutherland

**Author notes:** Corresponding author., 07455 727337.

## Abstract

Culverts are often installed under busy roads to help a variety of animals, from small frogs to bears, safely cross roads that bisect their habitats. One of the first roadway culvert systems designed specifically for amphibian use in the United States was installed along Henry Street in Amherst, Massachusetts in 1987 to protect spotted salamanders (*Ambystoma maculatum*). These salamanders cross Henry Street during their annual migration to their breeding pools. In recent years, anecdotal evidence from volunteers monitoring the site suggested that salamanders were no longer using the tunnels. To evaluate this concern we conducted salamander counts in 2016, 2017, and 2018 to quantify tunnel use. In 2016, only 11% of observed salamanders used the tunnels– a substantial decrease from 68% in 1988, one year after their installation, when the tunnels were last evaluated. Subsequently, we implemented two tunnel modifications in an effort to increase tunnel usage above the established 2016 baseline. Unfortunately, neither retrofit was successful. Previous studies have demonstrated that salamanders prefer minimum tunnel apertures of >0.4 m, so it is likely that the 0.2 m apertures here are inadequate. This may create differential light and humidity inside and outside the tunnels that is recognized by the salamanders. While many studies have evaluated amphibian tunnel use in lab and field settings, ours is one of the first studies to have examined tunnel usage data long after initial installation. These long-term data are critical for evaluating what factors are necessary for maintaining tunnels over decades-long time scales.

## Introduction

Roads and highways can cause substantial complications for wildlife, including habitat fragmentation, subdivision of once contiguous populations, and road mortality (Forman and Alexander 1998). Road mortality is particularly high for reptiles and amphibians, which move across roads slowly, often in large numbers (Gibbs and Shriver 2005, Holderegger and Di Giulio 2010). Given that an estimated one million vertebrates are killed on roads per day in the United States, a number that is increasing due to human population growth and increased road density (Forman and Alexander 1998, Vos and Chardon 1998), finding approaches that facilitate safe road crossings for animals is an important challenge. To attempt to reduce the effects of road mortality on indigenous fauna, the use of barrier fences and wildlife tunnels is becoming widespread (Forman and Alexander 1998, Taylor and Goldingay 2003, Dodd et al. 2004, Beebee 2013). The effectiveness of tunnels has been tested experimentally in closed conditions where workers have been able to vary tunnel parameters (Woltz et al. 2008) and in the field with wild populations (Jackson and Tyning 1989, Ashley and Robinson 1996, Hels and Buchwald 2001, Taylor and Goldingay 2003, Dodd et al. 2004, Aresco 2005, Gibbs and Shriver 2005, Yanes et al. 2005, Patrick et al. 2010). These studies have demonstrated that the combination of barrier fences and tunnels can drastically decrease road mortality in amphibians and reptiles. Although these studies have helped clarify the issue of road mortality and the effectiveness of tunnel systems, there have been few studies evaluating the continued success of older tunnel systems years after their installation.

One of the first amphibian tunnel systems in the United States was built in 1987 along Henry Street in Amherst, Massachusetts, to protect spotted salamanders (*Ambystoma maculatum*) during their annual migration (Fig. 1) (Jackson and Tyning 1989). Jackson and Tyning (1989) quantified the fence and tunnel effectiveness for the Henry Street tunnel system in 1988, one year after installation. They found that a total of 68.4% of the total recorded salamanders crossed through the tunnels, with 75.9% of salamanders that reached the tunnel entrance crossing through the tunnels. Since the early 1990s, volunteers have monitored this site during the spring migration and have carried salamanders that either climbed the drift fences or refused to use the tunnels across Henry Street safely. Anecdotal evidence from these volunteers suggested that over the years salamanders decreased their use of the tunnels. Rather than use the tunnels, salamanders balk at tunnel entrances and attempt to find other ways across the street, often ending up on the road surface. To address these concerns, we measured the effectiveness of the Henry Street tunnels in 2016, 2017, and 2018.

**Figure 1.**
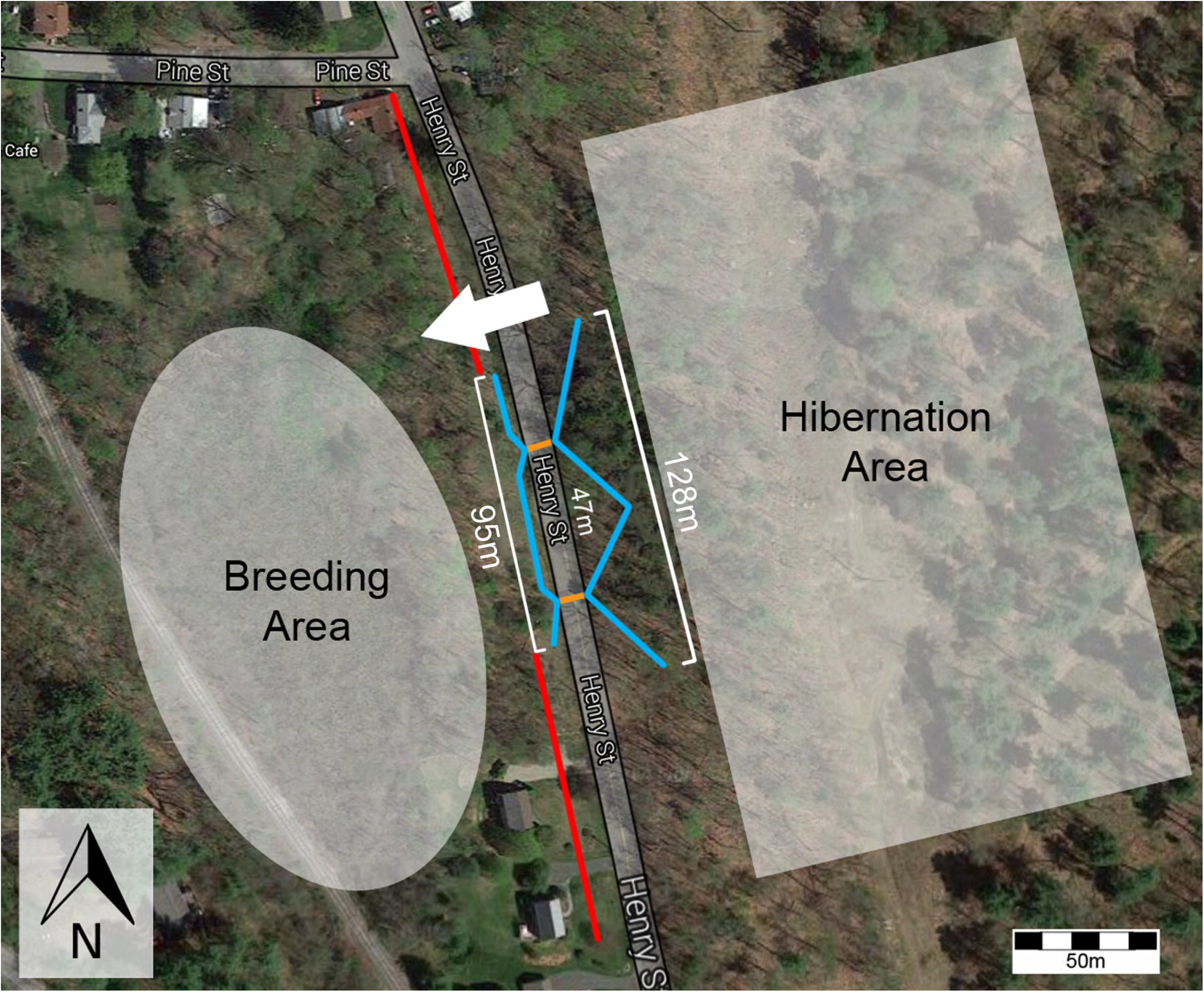
Site map of the spotted salamander road-crossing site. Henry Street bisects the salamander hibernation area and breeding area. The tunnels are marked in orange and the fences are marked in blue. The red lines show the full extent of the salamander crossing, although the vast majority of salamanders cross within the range of the fences.

Retrofitting tunnels may be a cost-effective alternative to reinstalling tunnels, which often exceeds town conservation project budgets. In 2016, we monitored the tunnels without performing any modifications in response to apparent decreased tunnel use. In 2017 and 2018, we tested two cost-efficient retrofits by experimentally manipulating one tunnel in each year, leaving the other as a control. In 2017, we investigated whether placing a light at the end of a tunnel would encourage use as suggested by Jackson (1996). In 2018, we constructed a platform leading down to the tunnel with a short drop-off just before the tunnel entrance to attempt to force salamanders to use the tunnels. Categorizing salamander count data into successful crosses, balks, and fence climbs, we estimated salamander mortality at this site and evaluated how current tunnel usage compares to usage just after tunnel installation in 1988 (Jackson and Tyning, 1989). In addition to improving the Henry Street wildlife tunnels, we sought to determine more generally if it is possible to retrofit older amphibian tunnel systems in a cost-effective manner.

## Methods

Spotted salamanders move in a mass migration from their hibernation areas to breeding pools once annually. Spotted salamander counts were conducted only during the migration from the hibernation area to the breeding area, not as they returned to their hibernation areas from the breeding pools. While the majority of a population migrates in a single night, often called a ‘big night,’ there are a small number of individuals that migrate either before or after. Our analyses and assessments are based solely on ‘big night’ data collected on the 10th of March (2016), the 28th of March (2017), and the 29th of March (2018), although additional data for migration nights were also collected (Table S1). Methods were carried out under IACUC 2016-0016.

### Initial assessment of tunnel functionality in 2016

Each year of the study, the tunnels were prepared for the migration several weeks prior to the anticipated event to ensure consistency. Tunnels were cleared of obstructions, areas near the tunnels were raked, and trash was collected in the vicinity. Additionally, fences were checked for gaps and were repaired as necessary.

During each migration event, volunteer citizen scientists were given an orientation to the tunnels and the experiments run each year and then were asked to walk along the road to tally the number of salamanders that climbed the drift fences. We also monitored the tunnel entrances to record the behavior of salamanders that reached the tunnels. Salamanders were counted as either ‘on road’, ‘successfully passed through tunnel’, or ‘balked at tunnel’. Volunteers were asked to carry salamanders found on the road to the west side of the street near the breeding habitat to ensure that the salamanders were not crushed by cars. We note that road counts are conservative estimates because some salamanders were found dead on the road and other salamanders may have successfully crossed the road without being found. However, given the large volunteer force (n > 20), small patrol area, and small number of fatalities, we consider the estimate to be representative of the larger pattern of road crossing efforts. Fatalities were not considered since it was impossible in many cases to distinguish dead spotted salamanders from other amphibians. A salamander was considered a “balk” if it either crossed in front of the entrance rather than approached the tunnel, or if it entered the tunnel but subsequently turned around, exited, and walked at least 50 cm away from the entrance. Upon balking, the salamander was carried safely across the street so that it would not be double-counted. The 2016 data served as a ‘baseline’ measure of the effectiveness of the tunnels prior to modifications in 2017 and 2018.

### A light at the end of the tunnel in 2017

Anecdotal evidence from prior years suggested that adding a light to the far end of the salamander tunnels increased usage (Jackson 1996). To test that hypothesis, we placed a bright white-light LED lantern (Black Diamond Apollo Lantern, 200 lumens) in a transparent, watertight plastic bag at the western end of the experimental (north) tunnel in 2017. No light was placed at the south tunnel (control) and only flashlights with dim red lights were used to patrol for salamanders. The south tunnel was chosen randomly as a control. Observation and tallying methods followed those of 2016.

### Salamander platform in 2018

We constructed a platform to place in front of one entrance such that salamanders that approached the tunnel would drop into a shallow pit with an 18 cm tall ledge just in front of the tunnel (Fig. S1). Climbing out of the pit was difficult and was qualitatively considered more energetically costly than using the tunnel. The goal was to use the platform to discourage balking and encourage tunnel use. The platform was constructed of pine struts overlaid by a plastic mesh, which was covered with soil to mimic natural ground cover (Fig. S1). As in 2017, the south tunnel was once again chosen randomly as the control and was not modified. The platform was placed in front of the north tunnel. Observation and tallying methods followed those of previous years.

### Statistical analyses

To estimate the probability of tunnel crossing versus balking, we used a binomial general linear model (GLM) with individual responses as the binary variable (1 = crossed, 0 = balked) for salamanders that reached the tunnels. Using a series of five competing models (Year, Tunnel, Year + Tunnel, Year * Tunnel, and a null model), we examined whether the probability and type of tunnel use varied by year (2016, 2017, 2018) and tunnel (north, south). The Year * Tunnel model predicted that salamanders would prefer one tunnel over another in a specific year (i.e., that tunnel modifications increased tunnel usage). We compared our models using Akaike Information Criteria (AICc–Burnham and Anderson 2002) corrected for small sample size, where the lowest AICc score represents the model best supported by the data. AICc weights (AICω) were then calculated to determine relative model support where ΣAICω = 1. All analyses were carried out in the base stats package in R (R Core Development Team 2017).

## Results

In 2016 (n = 124), the tunnel success was 11.3% for all counted salamanders (including fence climbers) and 20.6% for salamanders who reached the tunnels (Fig. 2a, Table 1). In 2017 (n = 108), 13.9% of the salamanders recorded successfully used the tunnels in total and 21.1% of the salamanders that reached the tunnels used them successfully. 25% of salamanders used the lit tunnel while 14.8% of salamanders used the dark tunnel successfully (Fig. 2b, Table 1). Finally, in 2018 (n = 357) only 7% of salamanders used the tunnels in total while 8.9% of salamanders that reached the tunnels used them. Of 113 salamanders that fell in the pit past the platform, only 20 crossed through the tunnel (17.6%) (Fig. 2c, Table 1, Table S1). The salamanders readily walked off of the platform into the pit and balking occurred after entering the pit.

**Figure 2.**
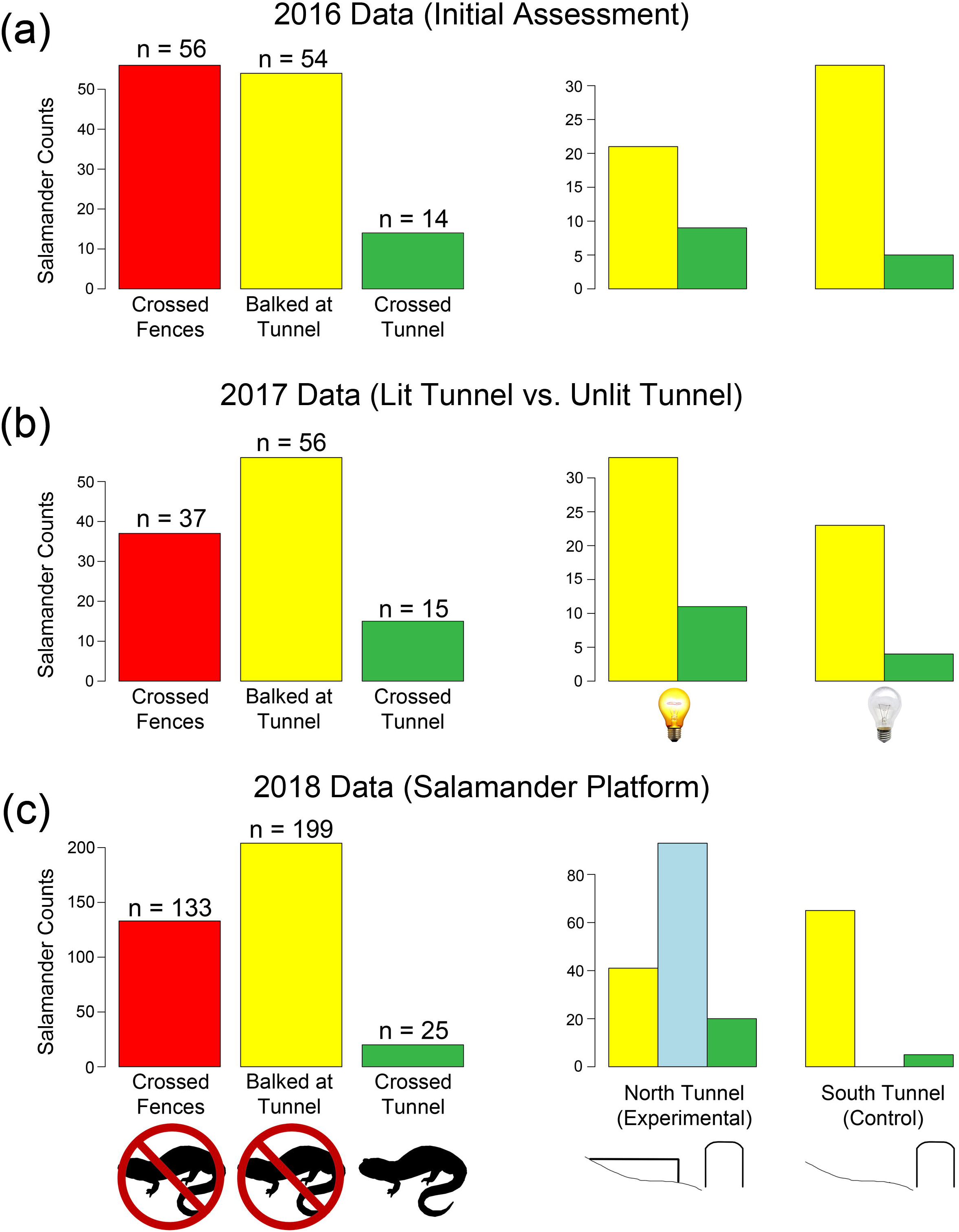
Salamander crossing data (2016–2018) showing the proportion of salamanders that crossed the fences prior to reaching the tunnels (red), salamanders that balked at the tunnel entrance (yellow), and salamanders that successfully used the tunnels (green). The northern and southern tunnel data are presented separately to show how well each tunnel performed (right). (**a**) 2016 data (control year). (**b**) 2017 data comparing the northern tunnel (lit tunnel) and the southern tunnel (control). (**c**) 2018 data comparing the northern tunnel (the salamander ramp) and the southern tunnel (control). For 2018, salamanders that fell into the pit following the ramp and climbed out are shown in blue while salamanders that approached the pit, but did not enter the pit are in yellow.

**Table 1.**
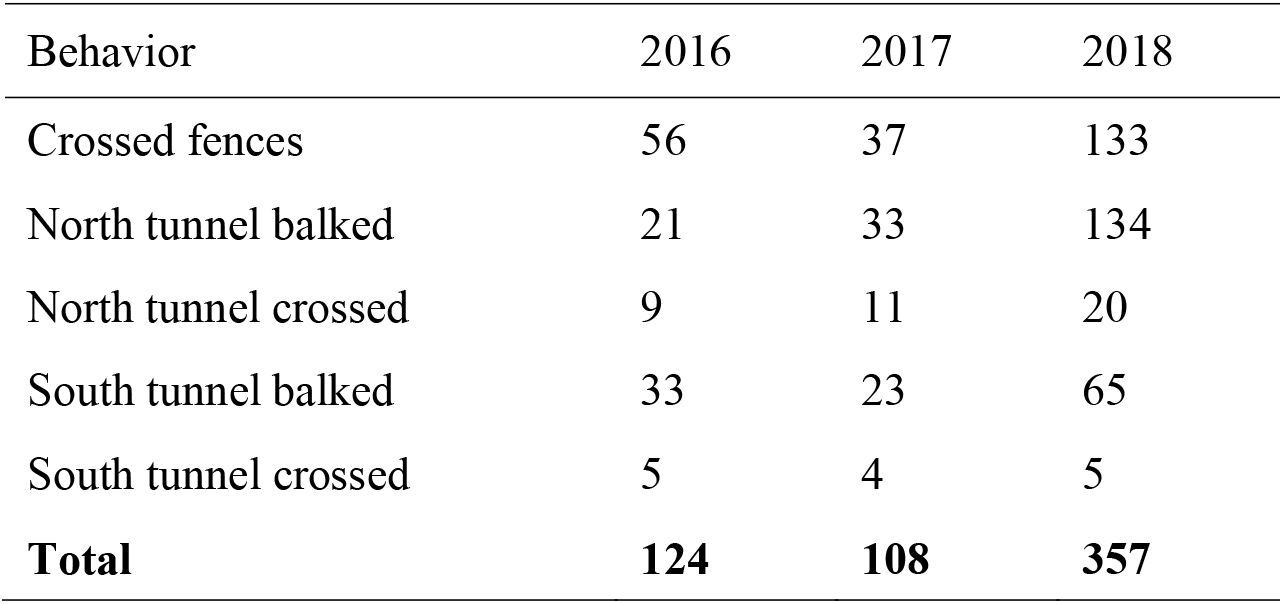
Raw counts of salamanders by tunnel and behavior for 2016–2018 big nights.

| Behavior | 2016 | 2017 | 2018 |
| --- | --- | --- | --- |
| Crossed fences | 56 | 37 | 133 |
| North tunnel balked | 21 | 33 | 134 |
| North tunnel crossed | 9 | 11 | 20 |
| South tunnel balked | 33 | 23 | 65 |
| South tunnel crossed | 5 | 4 | 5 |
| **Total** | **124** | **108** | **357** |

Comparing our models, there was compelling evidence that tunnel crossing success varied both by year and by tunnel (top three models in Table 2: Σ_1:3_ = 0.81). However, the model that allowed for variation by treatment effect (*Year * Tunnel*) was the lowest ranked model, receiving relative model support of just AICω = 0.05. This strongly suggests that there were negligible effects of both the installation of a light at the end of the tunnel and the addition of a platform. However, despite the large support for year and tunnel effects, there was still no clear top model and the top three models were all well supported (Table 2). To account for this uncertainty and to produce estimates of crossing probabilities for each year-tunnel combination, we generated a model-averaged prediction for each (Table 3). While the marginal effects of tunnel and year were not significant, retaining the tunnel and year effects separately was supported using AIC model selection (Table 2), and suggested that between-tunnel differences were larger than that of between-year differences. Although there was variability between the north and south tunnel and between years, salamander crossing probability did not exceed 20% for any tunnel or any year in spite of modifications (Table 3). We note that because we did not have replicates for either our light or ramp experiments given the presence of only two tunnels across three years, our statistical power for detecting small differences was limited. However, the overall trend of low tunnel usage in spite of attempted retrofits is very strong.

**Table 2.** AIC-ranks including all models, the number of parameters included in each model (K), the AIC*c* score (the *c* denotes that the correction for small samples was used), the differences between each model and the most supported model (∆AIC*c*), the AIC model weight which represents the relative support for each model (AICω), and finally, the cumulative model weights (Σ_1:j_).

| Hypothesis | $K$ | $AIC_c$ | $\Delta AIC_c$ | $AIC_w$ | $\Sigma_{1:j}$ |
| --- | --- | --- | --- | --- | --- |
| *Year + Tunnel* | 4 | 366.9 | 0.00 | 0.34 | 0.34 |
| *Tunnel* | 2 | 367.4 | 0.50 | 0.26 | 0.60 |
| *Year* | 3 | 367.8 | 0.96 | 0.21 | 0.81 |
| *Null* | 1 | 368.6 | 1.71 | 0.14 | 0.95 |
| *Year \* Tunnel* | 6 | 370.8 | 3.96 | 0.05 | 1.00 |

**Table 3.** Model averaged predictions of the probability that a salamander uses a given tunnel to cross the road.

| Year | Tunnel | Probability | SE |
| --- | --- | --- | --- |
| 2016 | North | 0.148 | 0.031 |
| 2017 | North | 0.190 | 0.050 |
| 2018 | North | 0.133 | 0.028 |
| 2016 | South | 0.110 | 0.033 |
| 2017 | South | 0.142 | 0.052 |
| 2018 | South | 0.097 | 0.031 |

## Discussion

When roads impinge upon amphibian habitats it is necessary to implement conservation measures to ensure that slow-moving amphibians are not locally extirpated. Amphibian tunnels or culverts in combination with drift fences have become a popular and successful method for mitigating the risks roads pose to wildlife (Gibbs and Shriver 2005, Yanes et al. 2005, Woltz et al. 2008). Older tunnels, such as those installed at Henry Street in 1987, should be carefully monitored over long timescales.

Although Jackson and Tyning (1989) found that the spotted salamanders used the Henry Street tunnels substantially more in 1988 (one year after installation) than during our study (30 years after installation), they did note some tunnel balking. Of 95 salamanders marked, 65 passed through the tunnels successfully (Jackson and Tyning, 1989). The balking was thought to potentially be a result of differences between the interior and exterior of the tunnels in temperature, humidity, illumination, airflow, human disturbance, or a combination of these factors (Jackson and Tyning, 1989). Jackson (1996) further suggested that a lack of light might lead to salamander hesitation and that the placement of a light at the far entrance of the salamander tunnels may increase use.

However, their data were not conclusive. The idea of increased light affecting tunnel use has been debated based on laboratory-based experiments where light permeability was shown to not be a significant factor affecting frog or turtle tunnel usage (Woltz et al. 2008). We found no substantial increase in tunnel usage when the north tunnel was experimentally lit in 2017 (Fig. 2b; Table 3) suggesting that illumination did not have a strong impact on salamander tunnel use at Henry Street. The platform modification to the north tunnel in 2018 was designed to force salamanders directly to the front of the tunnel entrance and discourage balking. However, there was no change in tunnel usage as a result of the platform (Fig. 2c; Table 3). Indeed, salamanders would enter the tunnel, travel less than half a meter into the tunnel, and then turn around. These salamanders then spent hours trying to climb out of the pit rather than use the tunnel that directed them toward their breeding area.

It is not clear which factors led to the decline in salamander use between 1988 (Jackson and Tyning, 1989) and 2016–2018. It is possible that monitoring populations too soon after wildlife tunnel construction may lead to inaccurate tunnel usage numbers and many authors call for long-term tunnel monitoring data (Glista et al. 2009, Beebee 2013). Based on previous findings and our results, we hypothesize two potential factors that may have led to a lack of tunnel usage at Henry Street: (1) tunnel aperture and (2) tunnel roof construction. Tunnel aperture is likely one of the most important variables in wildlife tunnel construction (Mata et al. 2008, Woltz et al. 2008). The Henry Street tunnel entrances are 0.2 meters wide and 0.25 meters tall. Several previous studies have found that reptiles and amphibians prefer tunnels with apertures greater than 0.4 meters (Woltz et al., 2008; Beebee, 2013). In contrast, Patrick et al. (2010) found that spotted salamanders did not choose tunnels on the basis of entrance aperture. However, the smallest aperture that they tested had a diameter of 0.3 meters, 50% wider than the Henry Street tunnels. As the present study was a field experiment, we were not able to vary tunnel aperture and thus were not able to assess with certainty that this was the cause of balking among salamanders at Henry Street. Tunnel roof construction is another critical parameter likely impacting tunnel usage (Woltz et al., 2008; Beebee, 2013). At Henry Street, there are small slots in the top of the tunnels that run the length of the tunnels measuring 1.5 cm in width and 6.5 cm in length, spaced 2.5 cm apart parallel to the road and 4 cm apart perpendicular to the road (Fig. S2). Although these slots allow enough moisture into the tunnels to ensure the substrate is wet, it is possible that the small size of these slots is affecting the relative humidity inside the tunnels (Jackson and Tyning, 1989). Future experimental studies should seek to confirm these hypotheses outside of a field setting to determine an acceptable moisture range and tunnel opening design for amphibians.

Fences play a critical role in wildlife tunnel systems (Cunnington et al. 2014). Aresco (2005) found 100% mortality in turtles at wildlife tunnels without fences. While Dodd et al. (2004) showed that after tunnel installation road kill counts dropped dramatically, animals such as tree frogs that were able to climb barrier fences were unaffected by the presence of the wildlife tunnels. In 1988, of 95 salamanders marked at Henry Street, 87 reached the tunnels. This suggests that only 8.4% climbed the fences that year (Jackson and Tyning 1989). In contrast, the percentage of observed salamanders climbing the fences was 45% in 2016, 34% in 2017, and 37% in 2018 (Table 1). Fences can quickly fall into disrepair and must be maintained (Baxter-Gilbert et al. 2013). It was evident that the Henry Street fences needed annual maintenance, with new holes and broken fence components found prior to the salamander migration every year. Although plastic mesh fencing was put in place to allow water to pass through the fences and reduce erosion (Jackson and Tyning 1988), the salamanders can easily climb the mesh using the perforations as toeholds. Additionally, the height of the fences (<20 cm tall in many sections) makes them relatively easy for the salamanders to climb. Fences taller than 0.6 meters (Beebee 2013) with a substantial overhang (Aresco 2005) would discourage climbing and improve the system.

Based on these data, we conclude that the Henry Street salamander tunnels are being used by a small percentage of observed salamanders. Attempts to improve the tunnels in a cost-effect manner proved ineffective. The small size of the tunnel apertures and the lack of adequately large perforations along the tunnel roofs may be creating a differential in moisture and light between the tunnels and the outside environment (Jackson, 1996). It is likely the large citizen science force that has been mobilized through the Hitchcock Center for the Environment, rather than the presence of the wildlife tunnels, has kept these salamanders from being locally extirpated (Sterrett et al., 2019). However, while volunteers can help salamanders cross streets, they only help in one direction because salamanders do not move *en masse* from their breeding area back to their hibernation area. Consequently, volunteers alone cannot prevent decline (Beebee 2013).

It is unknown if other older wildlife tunnel systems have similar issues to the Henry Street tunnels. We echo many workers who lament the lack of long-term data on amphibian tunnel use and call for more studies examining the effectiveness of wildlife tunnels after initial installation (Glista et al. 2009, Beebee 2013). Preferences in tunnel design appear to differ between taxa and there is no one solution to tunnel design that will work for all species (Lesbarrères et al., 2004). Additional studies will help to elucidate species-specific patterns so tunnel systems may be optimized for the specific taxon that they are meant to aid.

## Acknowledgements

We thank the many Hitchcock Center for the Environment volunteers who made this project possible, as well as assistance from the University of Massachusetts Wildlife Club and Shotokan Karate Club. We also thank David Munteanu (Clemson University), Zena Casteel (Cornell University), Kallin Lang, and Patricia Brennan (Mount Holyoke College) for help with fieldwork. We thank David Dunn for construction of the salamander platform. Finally, we thank Michael Forstner (Texas State University), Scott Jackson (UMass–Amherst), Jacob Kubel (Mass Wildlife), Arianne Messerman (University of Missouri), Reed Noss (Florida Institute for Conservation Science), Molly Grace (University of Central Florida), Rhett Rautsaw (Clemson University), and the Town of Amherst Conservation Commission for discussions regarding experimental set-up.

## Funding

This research was partially supported by the National Science Foundation Postdoctoral Research Fellowship in Biology (Grant #: 1612211 awarded to BPH) and the Town of Amherst.

Data availability statement: All data are included in the manuscript or in the supplemental information.

## Supplementary data

Supplemental Table 1: Salamander count data for migration nights with the greatest number of salamanders and additional smaller migration nights. Only the “big night” data were used in these analyses since additional nights lacked the volunteer numbers to collect accurate road monitoring data.

Supplemental Figure 1: (**a**) Northern tunnel eastern entrance. (**b**) Salamander ramp with drop-off installed in front of northern tunnel entrance. (**c**) Salamander ramp covered with soil as was in place during the migration event in 2018. Scale = 0.5 meters.

Supplemental Figure 2: Tunnel roof along the road showing the spacing of slots.

