## Supplementary figures and images for "Keeping salamanders off the streets: Evaluating one of the first US amphibian road tunnels 30 years later"

### Fig. S1

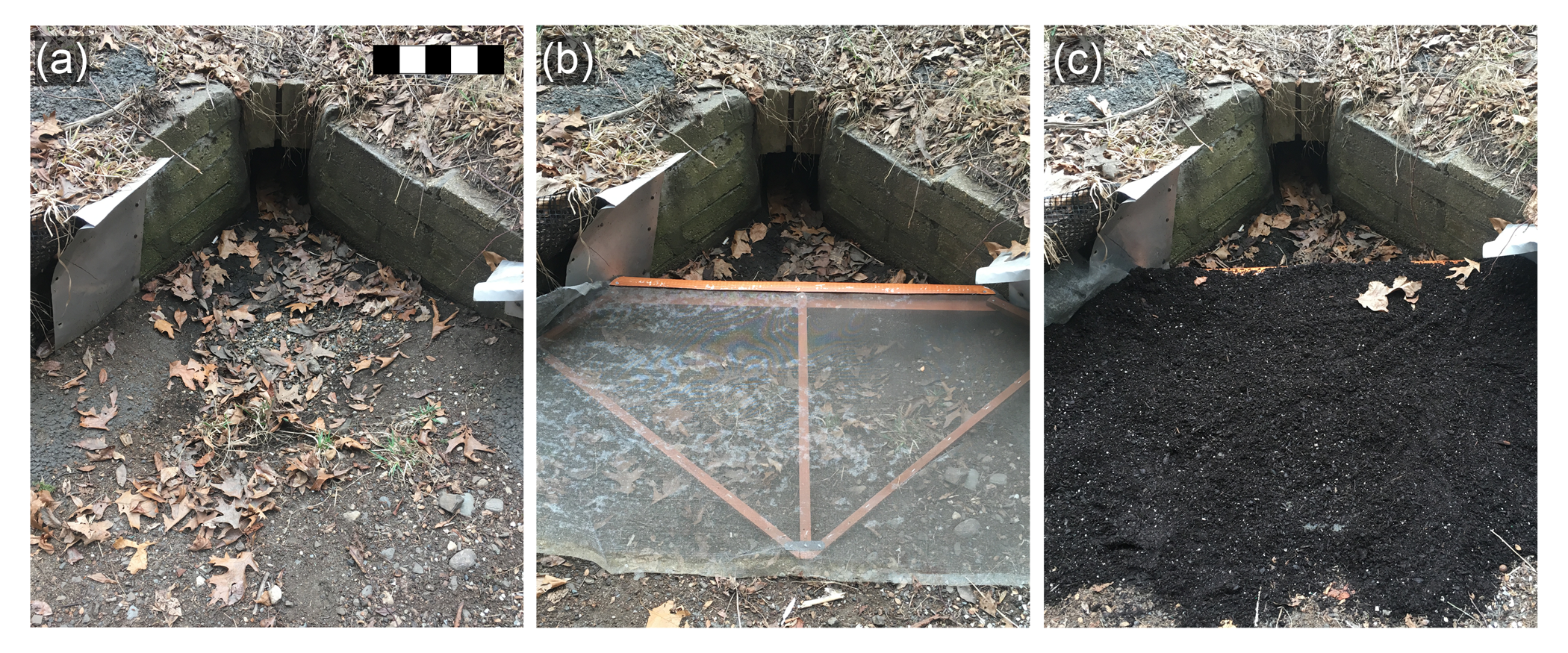

### Fig. S2

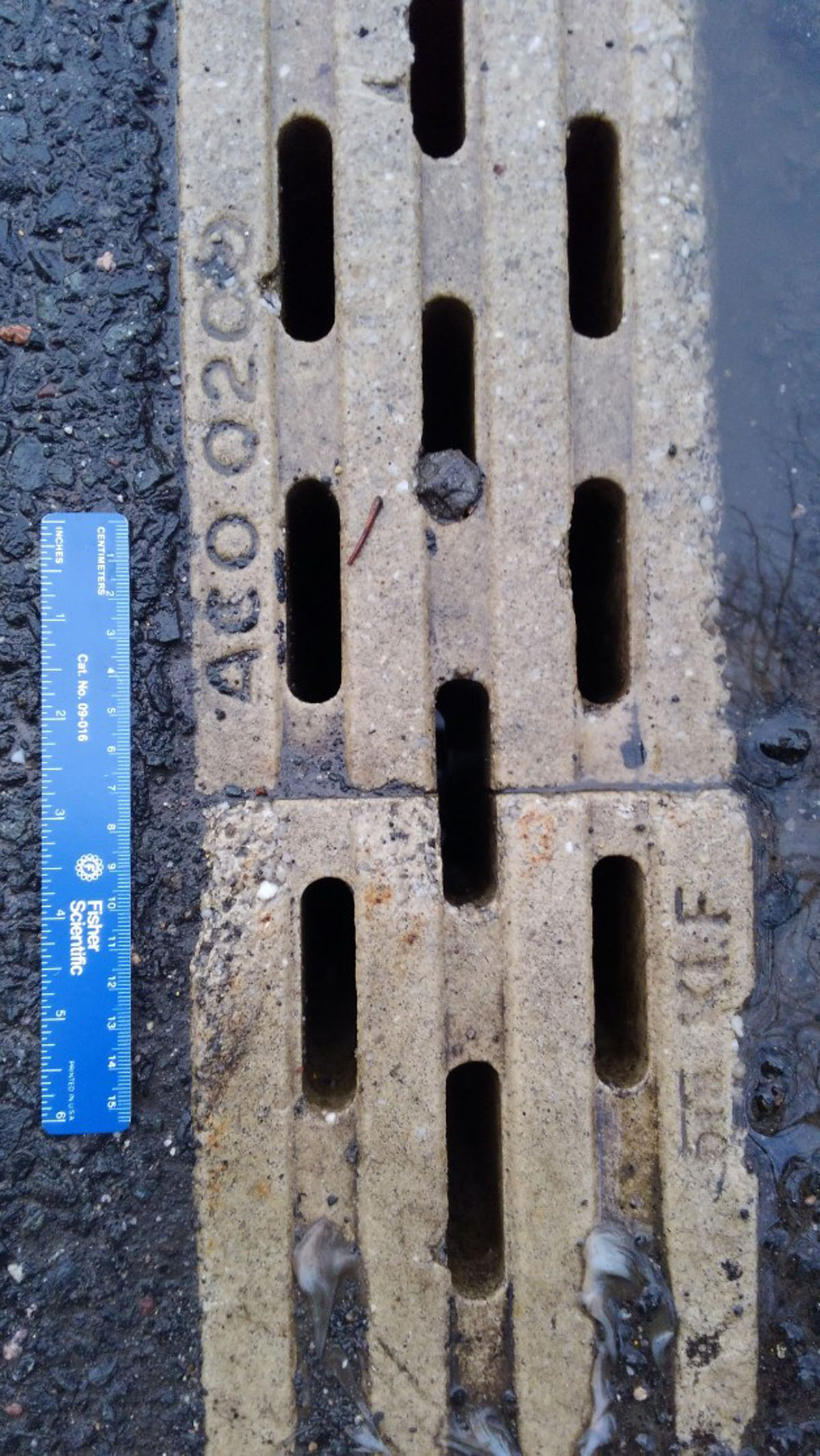
